# Genome-wide characterization of SARS-CoV-2 cytopathogenic proteins in the search of antiviral targets

**DOI:** 10.1101/2021.11.23.469747

**Authors:** Jiantao Zhang, Qi Li, Ruth S. Cruz Cosme, Volodymyr Gerzanich, Qiyi Tang, J. Marc Simard, Richard Y. Zhao

## Abstract

Therapeutic inhibition of critical viral functions is important for curtailing coronavirus disease-2019 (COVID-19). We sought to identify antiviral targets through genome-wide characterization of SARS-CoV-2 proteins that are crucial for viral pathogenesis and that cause harmful cytopathic effects. All twenty-nine viral proteins were tested in a fission yeast cell-based system using inducible gene expression. Twelve proteins including eight non-structural proteins (NSP1, NSP3, NSP4, NSP5, NSP6, NSP13, NSP14 and NSP15) and four accessory proteins (ORF3a, ORF6, ORF7a and ORF7b) were identified that altered cellular proliferation and integrity, and induced cell death. Cell death correlated with the activation of cellular oxidative stress. Of the twelve proteins, ORF3a was chosen for further study in mammalian cells. In human pulmonary and kidney epithelial cells, ORF3a induced cellular oxidative stress associated with apoptosis and necrosis, and caused activation of pro-inflammatory response with production of the cytokines TNF-α, IL-6, and IFN-β1, possibly through the activation of NF-κB. To further characterize the mechanism, we tested a natural ORF3a Beta variant, Q57H, and a mutant with deletion of the highly conserved residue, ΔG188. Compared to wild type ORF3a, the ΔG188 variant yielded more robust activation of cellular oxidative stress, cell death, and innate immune response. Since cellular oxidative stress and inflammation contribute to cell death and tissue damage linked to the severity of COVID-19, our findings suggest that ORF3a is a promising, novel therapeutic target against COVID-19.

**Significance:** The ongoing SARS-CoV-2 pandemic has claimed over 5 million lives with more than 250 million people infected world-wide. While vaccines are effective, the emergence of new viral variants could jeopardize vaccine protection. Antiviral drugs provide an alternative to battle against COVID-19. Our goal was to identify viral therapeutic targets that can be used in antiviral drug discovery. Utilizing a genome-wide functional analysis in a fission yeast cell-based system, we identified twelve viral candidates, including ORF3a, which cause cellular oxidative stress, inflammation and apoptosis and necrosis that contribute to COVID-19. Our findings indicate that antiviral agents targeting ORF3a could greatly impact COVID-19.

## Introduction

The ongoing SARS-CoV-2 pandemic due to coronavirus disease-2019 (COVID-19) is unprecedented in its rapid spread, persistence, and high fatality. Although current vaccines are effective in preventing SARS-CoV-2 infection, breakthrough infections are not uncommon. Moreover, the emergence of new viral variants could undermine the effectiveness of current vaccines. Antiviral drugs are an alternative, potentially effective means to curtail COVID-19 but currently such drugs are limited. The discovery of new antiviral agents is hampered by limited knowledge of the pathogenicity of the virus and of which viral protein(s) should be targeted. There is an urgent need to identify the viral proteins that cause harmful cytopathic effects, tissue damage, and cytokine storm, in order to advance the rational design of anti-COVID-19 therapies.

SARS-CoV-2 belongs to sarbecoviruses, a subgenus of β-coronaviridae family (1). It is one of the 7 human coronaviruses (hCoVs) that cause human diseases that range from the common cold to SARS (severe acute respiratory syndrome) and MERS (Middle East respiratory syndrome). Like other hCoVs, SARS-CoV-2 is an enveloped, positive-sense (+), single-stranded RNA virus with an average genome size of 29.7kb. Upon viral entry into a host cell, the (+)ssRNA viral genome is released into the cytoplasm where the two overlapping open reading frames 1a and 1b (ORF1a and ORF1b) are translated to produce two polyproteins, which are further processed by viral proteases to generate a total of 29 SARS-CoV-2 proteins including 4 structural proteins, spike (S), envelope (E), membrane (M) and nucleocapsid (N), 16 nonstructural proteins (NSP1 - NSP16), and 9 accessary ORFs (3a, 3b, 7a, 7b, 8, 9b,9c and 10) (2). Among them, ORF3a, ORF8, ORF9c and ORF10 are unique to SARS-CoV-2 (3).

We carried out a genome-wide functional screening of CoV-2 proteins to identify viral targets that can be used for antiviral drug discovery and testing. We reasoned that SARS-CoV-2 protein therapeutic targets: i) should be essential for virus survival; ii) should contribute to virus pathogenesis; iii) should confer measurable cytopathic effects that could be used as endpoints for high-throughput drug screening; iv) could be either an established or a novel antiviral drug target. We adapted a unique integrated approach, *viz*., a well-established fission yeast cell-based system to conduct genome-wide and functional analyses of the viral proteins, with the goal of identifying proteins that cause harmful cytopathic effects. Fission yeast (*Schizosaccharomyces pombe*) is a haploid single-cell eukaryotic organism that has been used extensively as a model to study human cancer biology (4) and viruses (5–8). Fission yeast is a well-tested model system for the study of highly conserved cellular activities (9, 10) such as the cytopathic effects caused by viral proteins, including inhibition of cell proliferation, interruption of cell structural integrity, and induction of apoptosis and necrosis (5). Fission yeast cell-based systems also have been shown to be suitable for antiviral drug testing and drug discovery (6, 11).

A total of 29 viral ORFs (3) representing the entire viral genome were cloned into the fission yeast gene expression system (7, 12). Expression of each viral ORF was under the control of an inducible *nmt*1 promoter (12), allowing all SARS-CoV-2-specific viral protein effects to be measured specifically by gene induction *vs*. without gene induction (5, 7). Using this unique integrated strategy, we report the identification of multiple possible viral therapeutic targets with a detailed description, as a proof-of-concept, of ORF3a. We demonstrate the potential of ORF3a to serve as a therapeutic target for the discovery and testing of antiviral drugs against COVID-19.

## Results

### Identification of SARS-CoV-2 cytopathic proteins that prevent cellular proliferation and colony formation of fission yeast

A total of 29 viral ORFs representing all viral proteins produced by the SARS-CoV-2 genome were expressed using inducible *nmt*1 promoter-mediated transcription in a wild type fission yeast strain SP223. Fission yeast cells expressing an empty cloning plasmid pYZ1N were used as a cloning vector (Vec) control. The effect of the SARS-CoV-2 protein expression on the ability of fission yeast to form colonies, an indication of cell proliferation, was measured on minimal selective agar plates (11, 12). All cells with viral genes repressed (*gene*-off) formed normal size colonies on the agar plates (**Fig. 1A; Fig. S2A**). In contrast, 8 non-structural viral protein-producing fission yeast cells (NSP1, NSP3, NSP4, NSP5, NSP6, NSP13, NSP14 and NSP15) and 4 accessory proteins (ORF3a, ORF6, ORF7a and ORF7b) showed either complete inhibition or near complete inhibition of yeast colony formation on minimal selective agar plates (**Fig. 1A**). In each case, either no colony or very small colonies were observed under viral gene-inducing conditions, consistent with cellular growth inhibition. With the Vec control, normal size colonies were observed on both gene-suppressing (*gene*-off) and gene-inducing (*gene*-on) plates, confirming the specificity of the viral protein effect observed. We named the 12 viral proteins the “cytopathic proteins”. Production of the remaining 18 SARS-CoV-2 proteins showed either no effect or reduced effect on the fission yeast colony formation (**Fig. S2A**).

**Figure 1.**
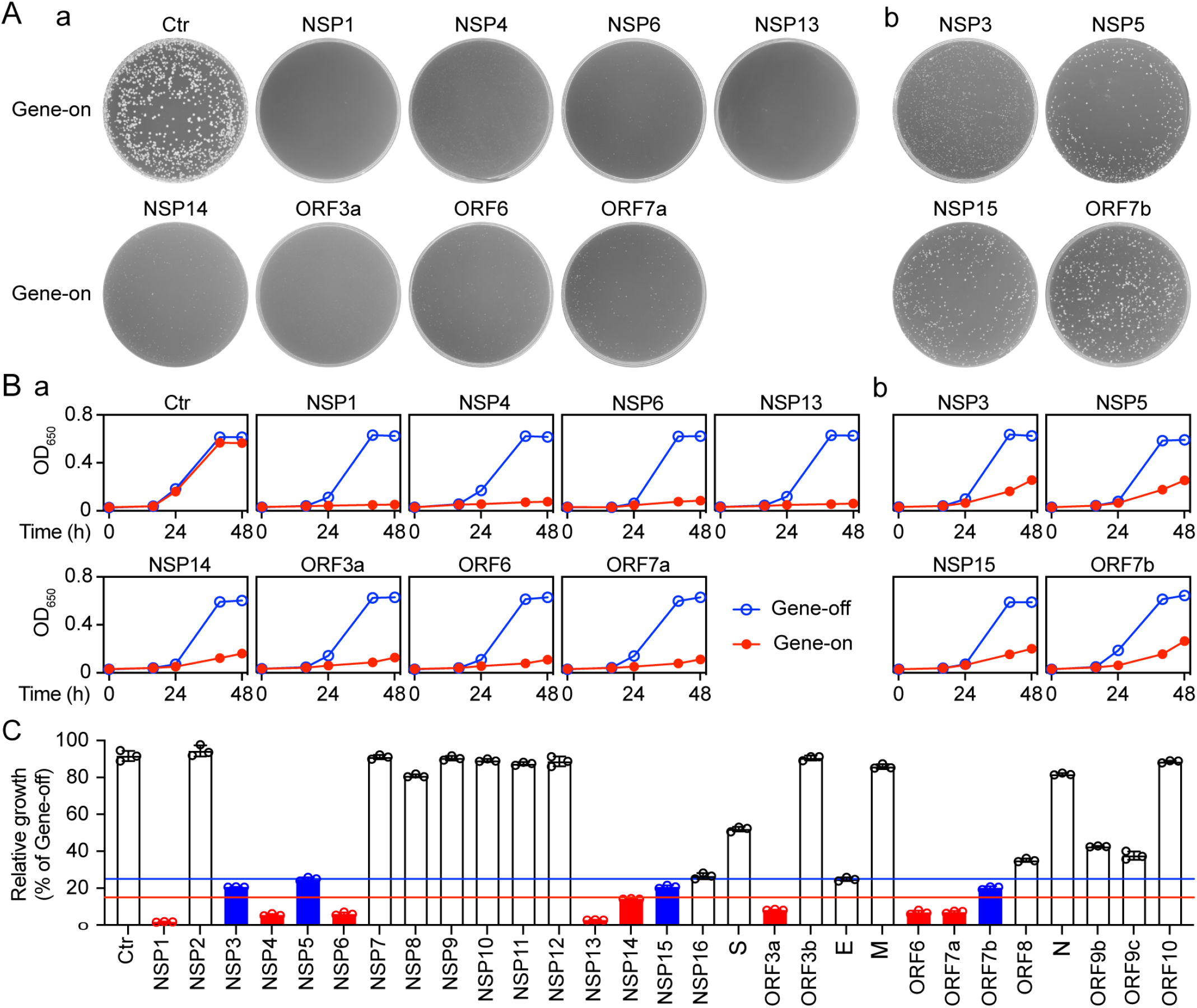
The effect of SARS-CoV-2 protein on cell proliferation. Effect of SARS-CoV-2 expression on fission yeast colony formation. (**A**), cellular growth (**B**) and summary of the relative cellular growth (15%, red; 25%blue) of each of the SARS-CoV-2 protein-expressing cells (**C**). The name of each SARS-CoV-2 protein is labeled above each agar plate. An empty pYZ1N vector (Ctr) was used as a control. Gene-off, no SARS-CoV-2 protein production; gene-on, the specific SARS-CoV-2 protein production was induced by triggering the *nmt*1 promoter-mediated gene transcription. Fission yeast colony formation was measured by growing SARS-CoV-2 protein-expressing fission yeast cells on the selective EMM agar plates and incubated at 30°C for 3–5 days before the pictures were taken. Cell proliferation analysis was carried out by comparing cellular growth between the SARS-CoV-2 protein-producing cells and the SARS-CoV-2 protein-suppressing cells over time. Cell growth was measured by spectrophotometry (OD_650_). Only the effect of those SARS-CoV-2 proteins that showed complete (NSP1, NSP4, NSP6, NSP13, NSP14, ORF3a, ORF6 and ORF7a) or nearly complete (NSP3, NSP5 and NSP15) inhibition of yeast colony formation is shown in (**A**) and thereafter. Complete data on cell proliferation are included in **Figure S2**. Each experiment was repeated at least three times and the standard errors of each time point were calculated.

To verify the effect of the 12 cytopathic proteins on cell proliferation, we measured the growth kinetics of the SARS-CoV-2-carrying fission yeast cells. Fission yeast cells were grown under the *gene*-off and *gene*-on conditions in the liquid minimal and selective EMM medium. Cellular growth was measured by the change in cell concentration over 48 h after gene induction (*agi*) using optical density (OD_650_)-based measurements. In the Vec only control cultures, two indistinguishable growth curves with typical logarithmic kinetics were observed in the *gene*-off and *gene*-on pYZ1N-carrying yeast cells. In contrast, expression of NSP1, NSP4, NSP6, NSP13, ORF3a, ORF6, or ORF7a completely blocked cellular growth, whereas expression of NSP3, NSP5, NSP14, NSP15 or ORF7b significantly suppressed cellular growth (**Fig. 1B**). During the first 24 h *agi*, both the SARS-CoV-2 *gene*-off and *gene*-on cells grew at about the same rate, consistent with 16 h being required to fully produce a protein after induction of genes under control of the *nmt*1 promoter (13). However, thereafter, growth of cells expressing the 12 viral proteins stopped or slowed compared to that of cells without viral gene expression. Consistent with the data on colony formation, the production of the remaining 18 SARS-CoV-2 proteins showed either no effect or reduced effect on fission yeast cellular growth (**Fig. S2B**).

### SARS-CoV-2 cytopathic proteins induce fission yeast cell morphologic changes and hypertrophy

Changes in cell morphology such as hypertrophy are linked to SARS-CoV-2-induced cytopathic effects in human airway epithelial cells and myocytes (14). To determine whether the SARS-CoV-2 cytopathic proteins affect cell morphogenesis, we compared microscopic cell morphologies between fission yeast cells with or without viral protein production. Control cells containing the Vec-carrying plasmid appeared normal under both gene-suppressing and gene-inducing conditions. Cells were typical rod-shaped with diameters of 3–4 μM and lengths of 7–14 μM. They were shiny on the edges, a sign of healthy cells (**Fig. 2A**) (9). Normal cell morphologies also were observed in cells that carried the 12 cytopathic protein-encoding plasmids under the gene-suppressing conditions. In contrast, expression of the 12 SARS-CoV-2 cytopathic proteins under the gene-inducing conditions resulted in various degrees of hypertrophic morphologies (**Fig. 2A**, bottom rows of each panel). The common features of the abnormal cell morphologies included cell elongation and enlargement. We further analyzed the overall changes in cell morphology caused by the 12 cytopathic proteins. Both the forward-scattered light (FSC) and side-scattered light (SSC) were measured for each cell population of 10,000 cells using flow cytometric analysis (**Fig. 2B**). The FSC is relative to the cell surface area, and thus measures cell size. The SSC is related to cell granularity, and thus determines intracellular complexity. Combined analysis of FSC and SSC provide an overall architectures of cell shapes in a heterogeneous cell population. As shown in **Fig. 2B**, there were significant differences in the cells’ overall architectures between the *gene*-off and *gene*-on cultures of cells with cytopathic proteins. In contrast, there were no clear differences between the gene-inducing and gene-repressing Vec-carrying fission yeast cells, suggesting the observed morphological changes were SARS-CoV-2 protein-specific. The production of all other SARS-CoV-2 proteins did not induce clear cellular hypertrophy (**Fig. S3B**). Together, our observations suggested that the 12 cytopathic proteins cause various degrees of cellular hypertrophy.

**Figure 2.**
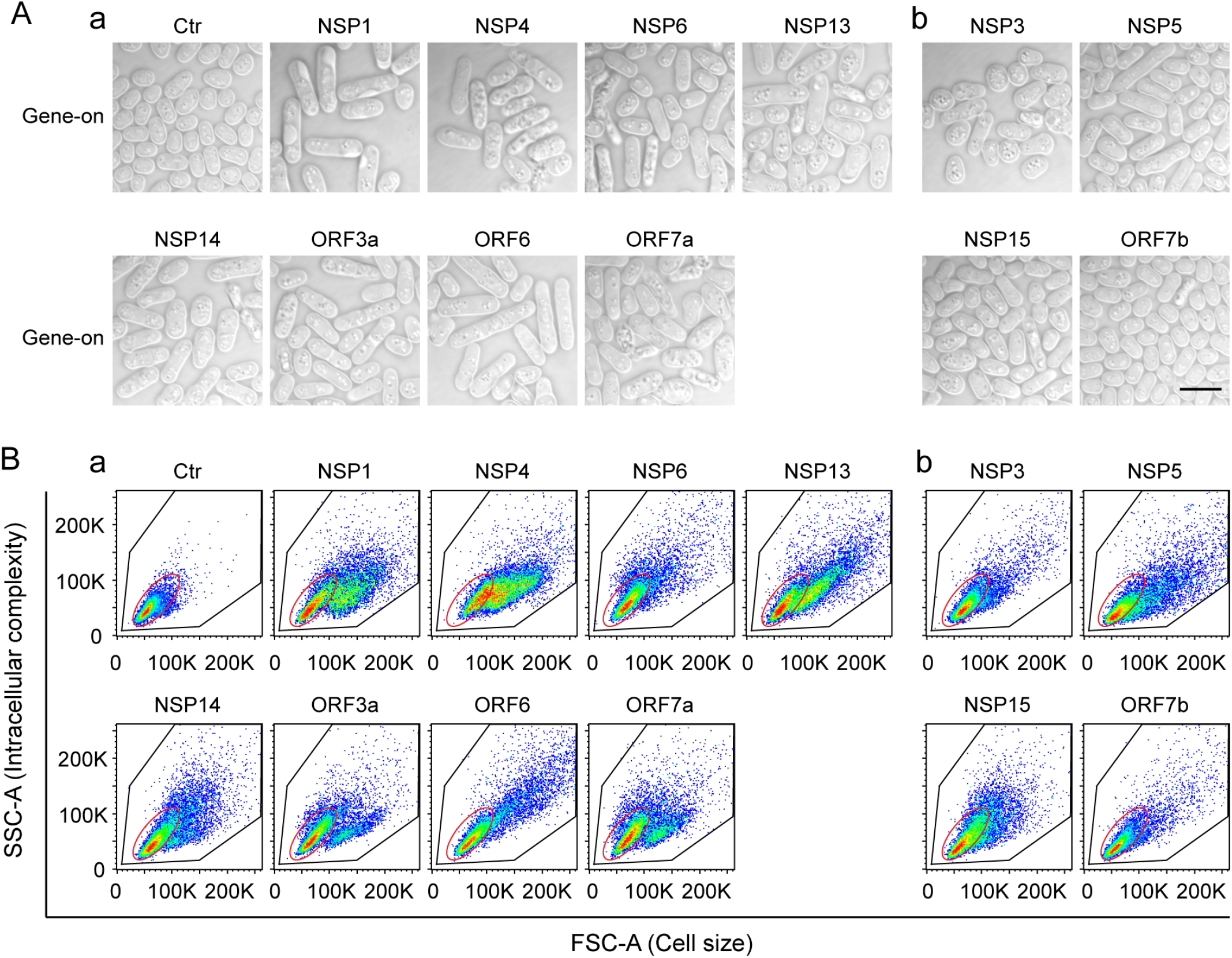
The effect of SARS-CoV-2 protein on fission yeast cellular morphology. Only those SARS-CoV-2 proteins that affected cell proliferation presented in **Figure 1** are shown here. Complete SARS-CoV-2 genome-wide data on fission yeast cellular morphology are included in **Fig. S3**. (**A**) shows the effect of individual SARS-CoV-2 proteins on fission yeast cell morphology. Each image was taken 48 h *agi* using bright field microscopy. Scale bar = 10 μM. (**B**) Overall cell morphology as shown by the forward scattered analysis. Ten thousand cells were measured 48 h *agi*. The forward-scatter (FSC) measures the distribution of all cell sizes. The side-scatter (SSC) determines intracellular complexity. Gene-off, no SARS-CoV-2 protein production; gene-on, SARS-CoV-2 protein produced.

### Correlation of SARS-COV-2 protein-induced cell death and induction of cellular oxidative stress in fission yeast

Since all 12 cytopathic proteins reduce or prevent cell proliferation (**Fig. 1**) and cause cellular morphologic changes (**Fig. 2**), these effects could be lethal. We tested whether the cytopathic proteins cause cell death. Inducible production of each viral protein was carried out as described above. Forty-eight hours *agi*, cell death was detected by Trypan blue, a vital diazo dye that specifically detects dead cells (5, 7). As shown in **Fig. 3A**, all cytopathic proteins induced cell death in the range of 10–60% (**Fig. 3C**, blue bars). In contrast, most of the other SARS-CoV-2 proteins did not kill cells (**Fig. S4A**).

**Figure 3.**
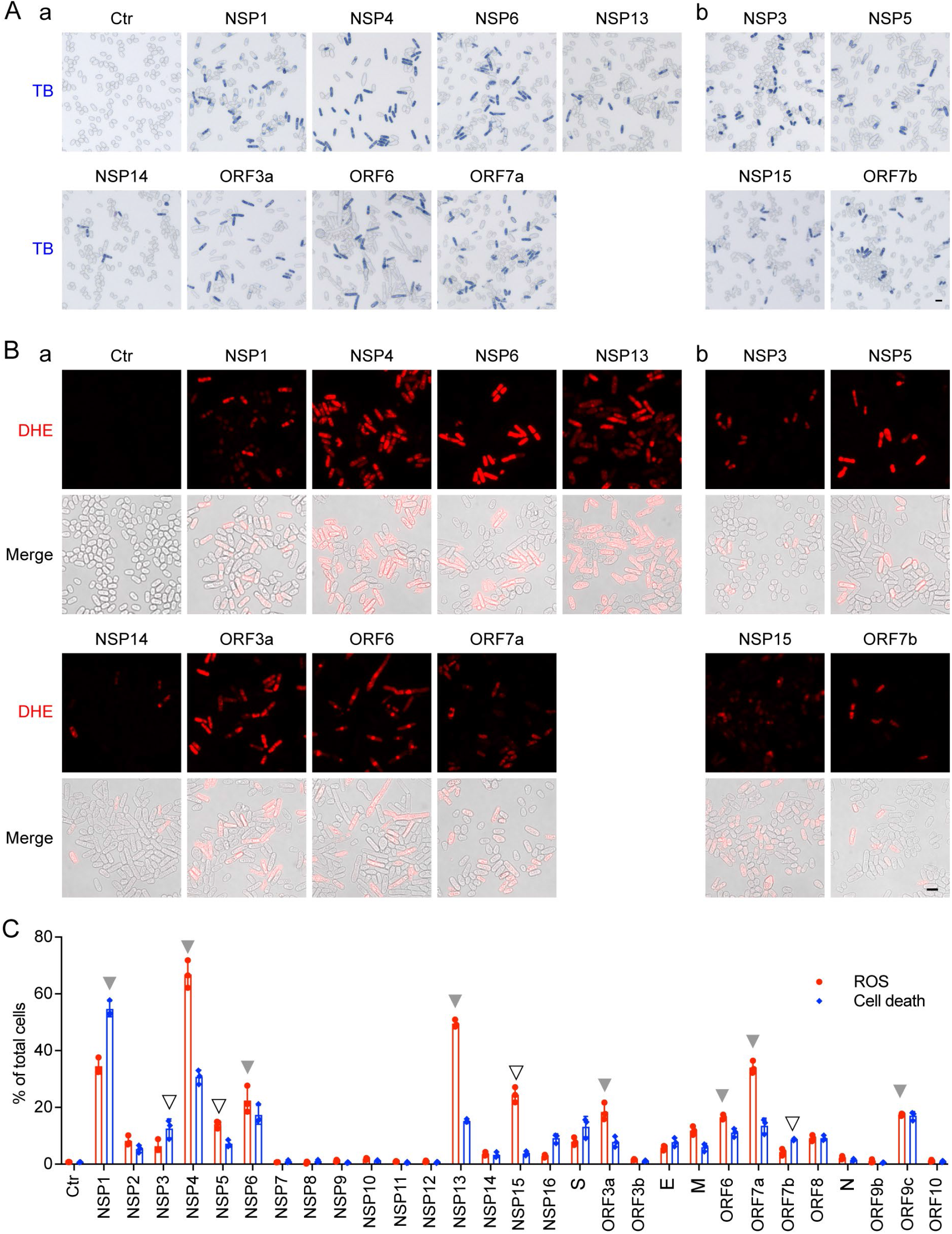
Correlation of SARS-CoV-2 protein-mediated cell death with induction of oxidative stress in fission yeast. (**A**) SARS-CoV-2 protein induces oxidative stress, as indicated by the DHE staining showing the production of ROS. Images were taken 48 h *agi*. Scale bar = 10 μM. BF, bright field; ROS, reactive oxidative species. DHE, an oxidative stress-specific dye (5, 58). (**B**) SARS-CoV-2 protein-induced cell death was measured 48 h *agi* by the trypan blue staining. (**C**) Quantitative correlation of SARS-CoV-2 protein-induced cell death (blue bars) and the production of ROS (red bars). Data represents mean ± SE from three independent experiments. Complete SARS-CoV-2 genome-wide data on fission yeast cell death and ROS production are included in **Fig. S4**.

We explored the mechanism of cell death caused by the cytopathic proteins. Because a cellular oxidative stress response can trigger cell death and is known to play an important role in COVID-19 (15), we tested for intracellular oxidative stress by measuring the production of reactive oxygen species (ROS). A ROS-specific dye, dihydroethidium (DHE) that produces red fluorescence in the presence of ROS, was used to measure the presence of ROS 48 h *agi* (**Fig. 3B**). Except NSP14, other cytopathic proteins triggered the induction of ROS as quantified and shown by the red bars (**Fig. 3C**). These observations suggest that cell death induced by the cytopathic proteins, with the exception of NSP14, was at least partially attributable to the induction of intracellular oxidative stress.

### ORF3a-induced apoptosis and necrosis are associated with the activation of mammalian cellular oxidative stress and immune pro-inflammatory responses

Of the identified cytopathic proteins, ORF3a was chosen for further study because it plays an important role in viral pathogenesis and its activities are linked to lung tissue damage and a cytokine storm in COVID-19 (16–18). Since we found that ORF3a-induced cell death was associated with the induction of cellular oxidative stress in fission yeast (**Fig. 3**), we tested whether a similar ORF3a effect is produced in mammalian cells, including human pulmonary epithelial A549, Calu-3 and kidney epithelial 293T cell lines. After transfection of the *ORF3a*-carrying plasmid into A549 and 293T cells, cell growth over time was measured, and cell viability and cell death were evaluated by the MTT method and trypan blue exclusion. As shown in **Fig. 4A-B**, the expression of *ORF3a* in either the A549 or the 293T cells significantly reduced cell growth, decreased cell viability, and increased cell death in comparison with the mock and the vector controls (Ctr). Possible effects of ORF3a-induced cell death *via* apoptosis or necrosis were evaluated by a RealTime Glo Annexin V Apoptosis and Necrosis assay. As shown in **Fig. 4C-a-b**, ORF3a-induced cell death involved both necrosis and apoptosis. ORF3a-induced apoptosis was mediated at least in part through the cleavage of caspase-3 (**Fig. 4C-c**). In addition, the expression of *ORF3a* triggered a cellular oxidative stress response, as measured by DHE staining (**Fig. 4D**). Our results suggest that ORF3a-induced apoptosis and necrosis in mammalian cells are associated at least in part with the induction of a cellular oxidative stress response.

**Figure 4.**
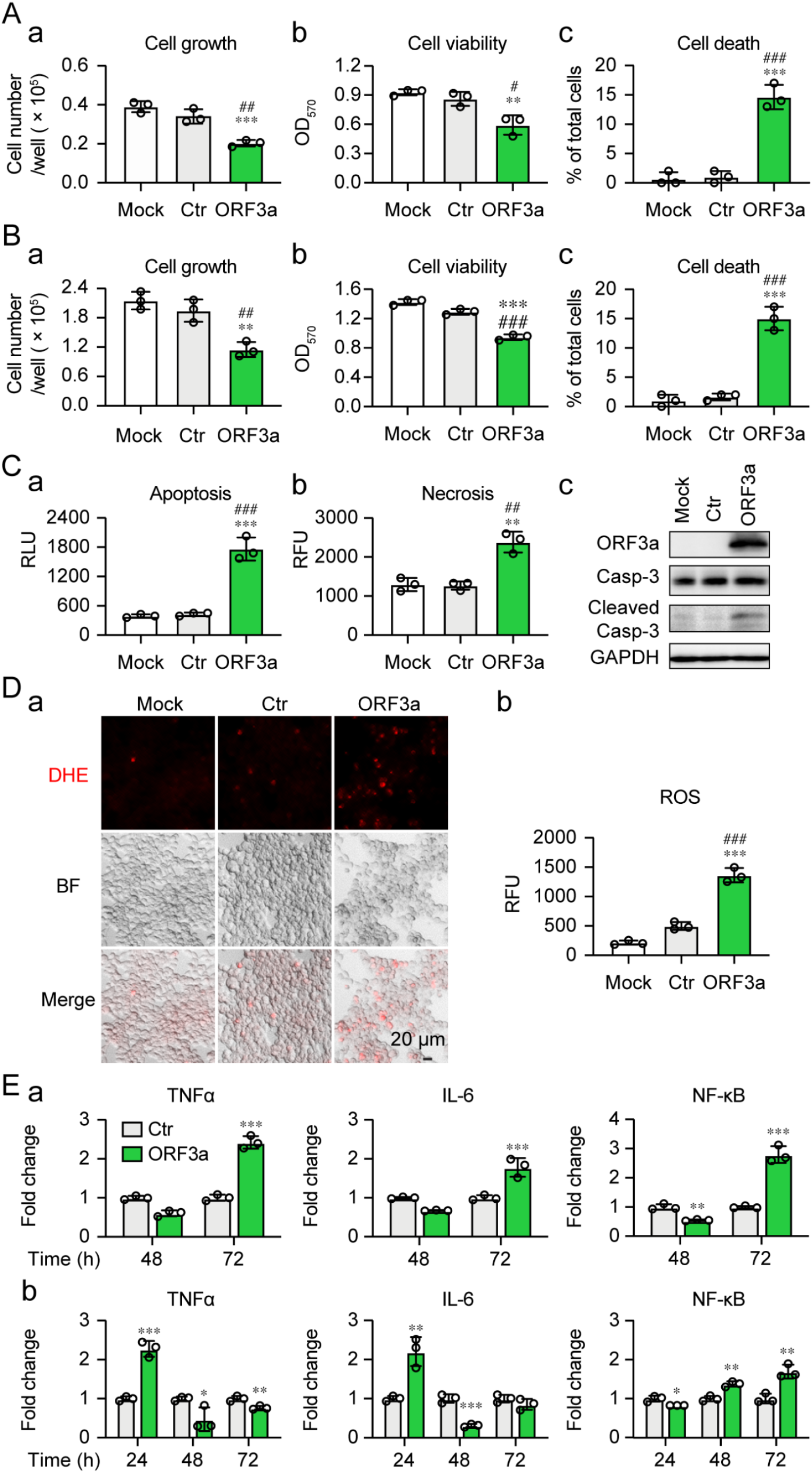
SARS-CoV-2 ORF3a-induced apoptosis and necrosis are correlated with the induction of cellular oxidative stress and innate immune pro-inflammatory responses in mammalian cells. Expression of SARS-CoV-2 *ORF3a* induces cellular growth reduction and cell death 72 *hpt* in human lung epithelial A549 cells (**A**) and human kidney epithelial 293T cells (**B**). (**C**) ORF3a induces apoptosis and necrosis 48 *hpt* measured by Annexin V (**a**), necrosis (**b**) and caspase-3 cleavage (**c**). (**D**) ORF3a triggers the induction of oxidative stress 48 *hpt* measured by the DHE straining. The *ORF3a* was cloned in a lentiviral constitutive expression vector (3). Scale bar = 20 μM. (**E**) ORF3a triggers elevated production of TNF-α, IL-6 and NF-κB in Calu-3 (**a**) and 293T (**b**) cells. Data are presented as mean ± SE from three independent experiments. Statistical differences between ORF3a and mock (indicated with #) or empty vector control (indicated with *) were evaluated. * or #, p < 0.05; ** or ##, p < 0.01; *** or ###, p < 0.001 (Pair-wise t-test).

To test whether the induction of apoptosis and necrosis by ORF3a are related to a host cellular immune response, we measured the production of the cytokines, tumor necrosis factor-alpha (TNF-α), interleukin-6 (IL-6) and their regulator, NF-κB (nuclear factor kappa B), by quantitative reverse transcriptase PCR (qRT-PCR). The mRNA levels of each gene target were measured over time (**Fig. 4E**). Gene transcription of both TNF-α and IL-6 increased moderately in Calu-3 and 293T cells 72 h post-transfection (*hpt*). The level of NF-κB also increased slightly over time. These data suggest that ORF3a-induced apoptosis and necrosis could also contribute to the activation of a cellular pro-inflammatory response.

### ORF3a-induced apoptosis and necrosis are affected by natural and artificial gene mutations

To further elucidate the molecular mechanism underlying ORF3a-induced apoptosis and necrosis (**Fig. 5**), we tested the mutational effect of ORF3a on the induction of apoptosis and necrosis. A natural mutant variant (Q57H) that is associated with the ongoing Beta variant, and deletion of a highly conserved guanine residue (ΔG188) were tested. The ΔG188 mutant was chosen because the amino acid (a.a.) 188 is highly conserved among sarbecoviruses, a subgenus of β-coronaviruses. G188 may be structurally important as it is one of the two guanidine resides that separate two antiparallel β4 and β5 sheets, and both residues are in the proximity of two homodimers of ORF3a and are within the inner cavity of the protein (19). The same methods as described above (**Fig. 4**) were used to measure cell growth, cell viability and apoptosis and necrosis with the ORF3a mutants. In comparison with wildtype ORF3a, the ΔG188 mutant showed significant differences in its effects including markedly reduced cellular growth, viability and increased cell death over time (**Fig. 5A-a-c**, red bars). In contract, the Q57H mutant showed slightly improved cellular growth, viability and reduced cell death (**Fig. 5A-a-c**, blue bars). Overall, the effects of the two ORF3a mutants on cellular growth and viability were inversely correlated with the effects on overall cell death (**Fig. 5A-c**), apoptosis and necrosis (**Fig. 5B**). Compared to wildtype ORF3a, the ΔG188 mutant markedly increased apoptosis and necrosis, whereas the Q57H mutant showed reduced apoptosis and necrosis. Consistent with the idea that ORF3a-induced cell death is mediated through the induction of cellular oxidative stress, a positive correlation also was seen between the wildtype and mutant ORF3a in the production of ROS (**Fig. 6**). Based on these observations, ORF3a-induced apoptosis and necrosis are likely mediated through the induction of cellular oxidative stress, and the overall structure of the ORF3a protein appears to be important for the induction of cell death.

**Figure 5.**
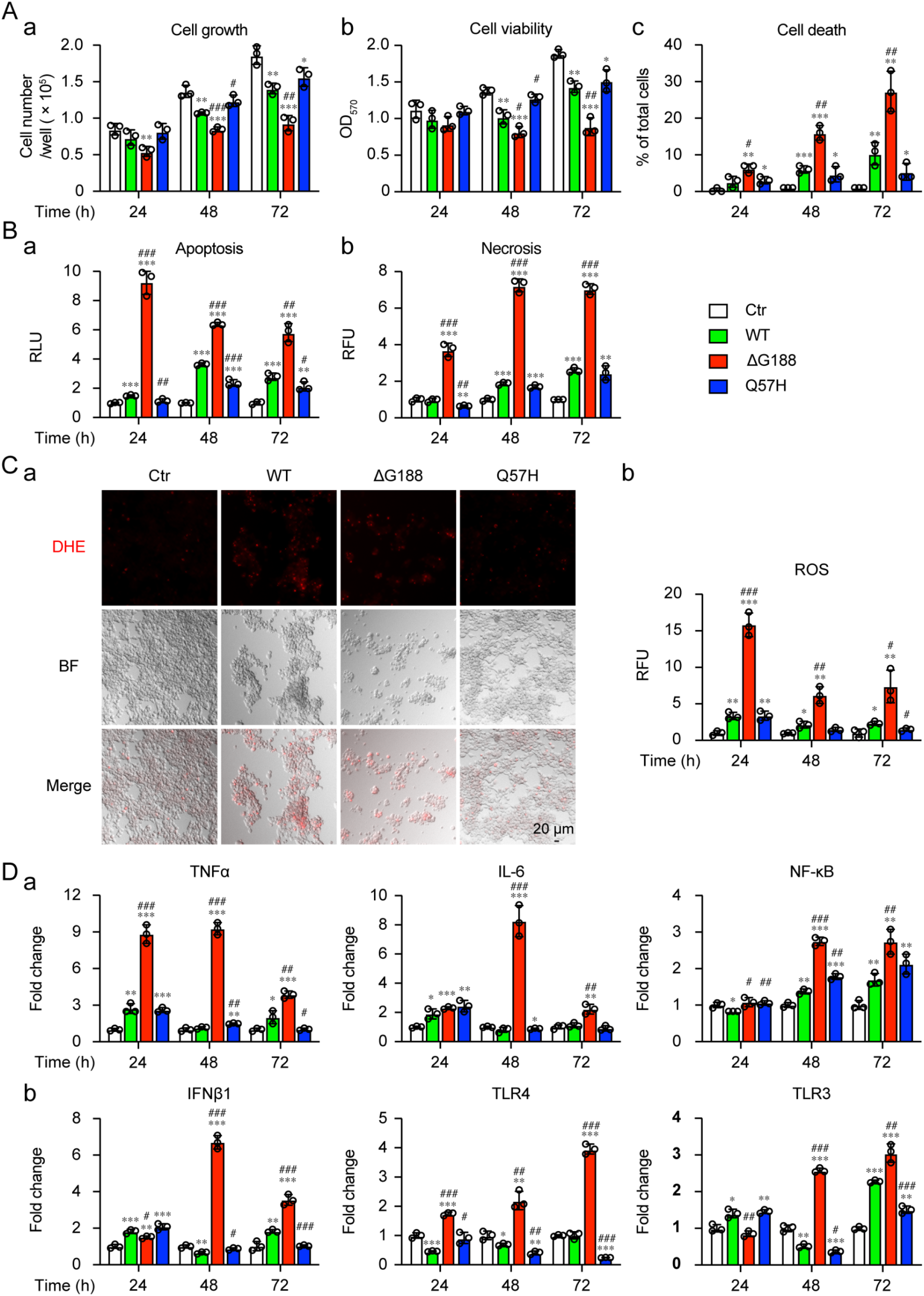
ORF3a-induced apoptosis by natural and artificial mutant variants correlate with cellular oxidative stress and innate immune pro-inflammatory responses. 293T cells were transfected with plasmids harboring ORF3a wild type (WT), ΔG188 or Q57H mutant variant. Effects of ORF3a mutant variants on cytopathic effects (**A**), as measured by cellular growth (**a**), cell viability (**b**) and cell death (**c**) at the indicated times. (**B**) Effects of ORF3a mutant variants on apoptosis (**a**) and necrosis (**b**). (**C**) Induction of oxidative stress. The images were taken at 72 *hpt*. Scale bar = 20 μM. (**D**) Activation of cellular innate immune pro-inflammatory responses, as measured by qRT-PCR. Data are presented as mean ± SE from three independent experiments. Statistical differences between control and ORF3a WT or mutants (indicated with #) or between WT and mutants (indicated with *) were evaluated. * or #, p < 0.05; ** or ##, p < 0.01; *** or ###, p < 0.001 (one-way ANOVA).

### ORF3a-induced apoptosis and necrosis are correlated with the level of cellular pro-inflammatory response

Since ORF3a-induced cell death is associated with the activation of host cell pro-inflammatory response (**Fig. 4E**), we tested whether the level of ORF3a-induced cell death is correlated with the level of the cellular pro-inflammatory response. Both Q57H and ΔG188 mutants were tested in comparison with the wildtype ORF3a. While the Q57H mutant showed a similar immune expression profile as the wildtype ORF3a (**Fig. 5D-a**, blue bars), sharp increases in TNF-α, IL-6, and NF-κB were observed with the ΔG188 mutant (**Fig. 5D-a**, red bars), suggesting a strong correlation between the level of apoptosis and necrosis induced by the ΔG188 mutant and the level of TNF-α, IL-6, or NF-κB. To further test whether other cytokines or pattern recognition receptors that recognize dsRNA virus also respond to the expression of ORF3a, we measure gene transcription of type I interferon beta-1 (IFN-β1), Toll-Like receptors 4 (TLR4) and Toll-Like receptors 3 (TLR3). Similar or slightly reduced activation of IFN-β1, TLR4 and TLR3 was observed in the Q57H mutant compared with the wildtype ORF3a (**Fig. 5D-b**), whereas sharp increases of IFN-β1, TLR4 and TLR3 were observed with the ΔG188 mutant (**Fig. 5D-b**). Interestingly, there were temporal differences in the activation of the various cytokines and regulators. The increase in TNF-α began as early as at 24 *hpt* and was significantly reduced at 72 *hpt* (**Fig. 5D-a**). In contrast, no significant increase of signals of NF-κB or TLR4 or TLR3 was seen at 24 *hpt*, but mRNA production gradually increased from 48 to 72 *hpt*, suggesting these might be late events. The two type I interferons, IFN-β1 and IL-6, showed a significant spike at 48 *hpt*. These observations suggest that there is an active interaction between the expression of ORF3a and the host cellular innate immune responses.

## Discussion

The goal of this study was to identify SARS-CoV-2 viral targets that can be used in high throughput drug screening and antiviral drug discovery. Our criteria for appropriate targets were that the viral protein must contribute to viral pathogenesis and cause measurable cytopathic effect that can be used in large-scale drug screening. Twelve viral proteins were identified that included 8 non-structural proteins (NSP1, NSP3, NSP4, NSP5, NSP6, NSP13, NSP14 and NSP15) and 4 accessory proteins (ORF3a, ORF6, ORF7a and ORF7b). Overall, they all inhibited cellular growth (**Fig. 1**), caused cellular hypertrophy (**Fig. 2**), and induced cell death (**Fig. 3**).

Two major types of viral proteins were identified, viral enzymes and viral proteins that participate in viral replication or transcription (20). Among the viral enzymes, NSP5 encodes the major 3C-like protease (3CL^pro^), NSP3 produces a minor papain-like protease (PL2^pro^), NSP13 encodes a viral helicase that unwinds duplex RNA or DNA with a 5’→3’ polarity, and NSP14 is a dual function enzyme with N-terminal exoribonuclease (ExoN) and C-terminal N7-methyltransferase (N7-MTase) activities. All these viral enzymes are highly conserved among coronaviruses and are essential for viral replication (20). Therefore, they might be ideal targets for antiviral drug discovery. Notably, viral proteases are proven antiviral targets against HIV-1 (21) and HCV infection (22). Some of the FDA-approved HIV-1 and HCV protease inhibitor drugs inhibit 3CL^pro^ or PL2^pro^ (23, 24). Although no clinical benefit has yet been shown from using those repurposed drugs, they nevertheless indicate the potential of using protease inhibitor drugs in treating COVID-19. The successful development of the first protease inhibitor antiviral drug Paxlovid (PF-07321332) against SARS-CoV-2 by Pfizer (25) attests this possibility. The fission yeast 3CL^pro^ or PL2^pro^ cell-based assays described here could conceivably be used for high-throughput screening and testing of future SARS-CoV-2 protease inhibitors. This is certainly feasible because our early study shows that all FDA-approved HIV-1 protease inhibitor drugs suppress HIV-1 protease in the fission yeast cell-based system (11). NSP13 also shows dNTPase and RNA 5’-triphosphatase activities besides its helicase activity. NSP13 is the most conserved coronaviral protein and it is essential for viral RNA transcription and replication. Therefore, it is another promising viral target for antiviral drug development (26).

Three of the identified viral candidates (NSP3, NSP4 and NSP6) form a core protein complex that is responsible for the formation of virus-induced double-membrane vesicles (DMVs). DMVs are part of the viral replication-transcription complex (RTC) in SARS-CoV-2 infected cells that drives replication and transcription for virus reproduction (27, 28). The RTC formed by NSP3/4/6 in SARS-CoVs links the cytoplasmic membrane with the endoplasmic reticulum lumen to induce membrane curvature and form unique DMVs (29). In addition, NSP3/4/6 interacts with E, M and N proteins to facilitate viral assembly (30). Thus, interruption of the NSP3/4/6 protein complex and DMV formation could potentially be a critical viral target to inhibit viral replication and transcription.

ORF3a was chosen for mechanistic studies because it fulfills the pre-defined criteria as a potential therapeutic target. ORF3a is a multifunctional protein that has 275 a.a. (~31 kD) and presents as a homodimer or tetramer (31). It has 3 transmembrane regions that span halfway across the membrane and cytosol that are linked to multifunctionalities including virulence, infectivity, ion channel formation, and virus release (16). ORF3a plays an important role in SARS-CoV-2-mediated viral pathogenesis, including induction of the inflammasome in damaged lung tissue, and initiation of cytokine storm (17, 18). High titers of anti-ORF3a antibody were found in SARS-CoV-2-infected patients, suggesting its clinic significance (32). Deletion or transcription knockdown of *ORF3a* from the SARS-CoV viral genome results in a major reduction of virus growth (33, 34). SARS-CoV-2 ORF3a is essential for viral replication in the absence of the E protein (34, 35).

Consistent with our findings from the fission yeast studies, ORF3a prevented cell proliferation and induced apoptosis and necrosis in mammalian cells (**Fig. 4A-C**), in part through the induction of cellular oxidative stress (**Fig. 4A-D**). ORF3a-induced apoptosis has been reported previously (36). However, oxidative stress-induced cell death had not been reported in SARS-CoV-2 infection, and the viral protein(s) responsible for inducing cellular oxidative stress had not been identified. Here, we report for the first time that ORF3a not only induces ROS production leading to cellular oxidative stress, but this cellular stress response appears to be a common feature of all the viral proteins except NSP14 that link to cell death in fission yeast cells (**Fig. 3C**, shown by arrows). Whether the activation of cellular oxidative stress also contributes to the death of human cells caused by other viral proteins besides ORF3a remains to be elucidated. Oxidative stress is a common cellular response to viral infection (37). Sustained viral infection leads to accumulation of the ROS in cells or tissues that may result in tissue damage and cell death (38, 39). Increasing evidence suggests that excessive ROS production is a major cause of local or systemic tissue damage and contributes to the severity of COVID-19 (40, 41). Therefore, targeting these viral targets could potentially help ease COVID-19. It is unclear at present why NSP14-induced cell death did not trigger a cellular oxidative stress response. One possible explanation is that it functions as an enzyme that is part of normal cellular activity.

ORF3a-induced apoptosis and necrosis might also be attributed in part to the activation of a cellular pro-inflammatory response, as three of the common pro-inflammatory cytokines, TNF-α, IL-6, and IFN-β1, were all elevated (**Fig. 4E**; **Fig. 5D**). Clinical studies suggest that high serum IL-6 and TNF-α levels are strong, independent predictors of patient survival (42, 43), and that hyper-inflammatory responses in patients with COVID-19 are a major cause of disease severity and death (42, 43). ORF3a-induced apoptosis and necrosis might be one of the viral factors that contributes to the elevation of IL-6 and TNF-α observed in these patients. Our data show that elevated production of IL-6 and TNF-α could be mediated through NF-κB, a well-recognized master regulator that mediates cellular pro-inflammatory responses to viral infection (44). In addition, TLR3 and TLR4 might also be involved in these elevations, as they are also activated (**Fig. 5D**). Both TLR3 and TLR4 recognize dsRNA viruses and trigger antiviral production of type I IFNs and proinflammatory cytokines. TLR4 is a cell membrane receptor (45) and TLR3 resides on the endosomal membrane in epithelial cells (46). TLR3 induces pro-inflammatory cytokine production through TRIF (46); whereas TLR4-mediated pro-inflammatory cytokine production is likely through NF-κB. Both TLR3 and TLR4-mediated pro-inflammatory responses contribute to the severity of the COVID-19. TLR4-mediated production of IL-6 and TNF-α is associated with the severity of COVID-19 in patients with cardiometabolic comorbidities (47). Inhibition of TLR3 by famotidine decreases IL-6 (48) and reduces the risk of intubation and death in patients hospitalized with COVID-19 and alleviates symptoms in non-hospitalized patients with COVID-19 (49). Both cellular inflammation and oxidative stress contribute to the severity of the COVID-19 (50). In addition, ORF3a-induced apoptosis and necrosis are a highly conserved cellular response among yeast (this study), *Drosophila* (51), and human cells (36). Therefore, targeting ORF3a-induced apoptosis and necrosis could be a promising antiviral strategy to fight against COVID-19. Fission yeast could potentially be used as a surrogate system for large-scale drug screening and testing of anti-ORF3a inhibitors.

The ΔG188 mutant triggered much stronger activation of the cellular oxidative stress and innate immune responses compared to wildtype ORF3a (**Fig. 4–5**). Early protein structural analysis suggests the G188 residue may be structurally important, as it is one of the two highly conserved guanidine resides among sarbecoviruses that separate two antiparallel β4 and β5 sheets, and both residues are in the proximity of two homodimers of ORF3a and are within the inner cavity of the protein (19). The ΔG188 mutant may interrupt the structure of the two antiparallel β4 and β5 sheets at the C-terminal end or the inner cavity where it affects ion channel activity or interaction of ORF3a with host cellular proteins. Structurally, the ORF3a activity has been linked to seven sequence motifs from the signal peptide (a.a. 1–15) of the N-terminus to a di-acidic motif (a.a. 171–173) (16, 36). This is the first demonstration of the functional relevance of the G188 residue at the β4/β5 junction of the C-terminus of ORF3a in the activation of cellular stress, innate immune response, and induction of apoptosis and necrosis. Additional mutagenesis studies are needed to confirm our findings. One possible explanation for the stronger effect of the ΔG188 mutant comparted to wildtype is that the wildtype ORF3a effect might be restricted by a host restriction cellular protein(s) through direct protein-protein interaction as shown in other viral infection (52). If this is indeed the case, interruption of the ORF3a protein with the host restriction factor(s) by the ΔG188 mutation could releases the ORF3a resulting enhanced triggering of host cellular stress and innate immune response leading to stronger cell death. This scenario is further suggested by the fact that the wildtype ORF3a completely kills fission yeast cells but only partially kills human cells. Note that ORF3a-induced cell killing in fission yeast is not an artifact caused by protein overproduction, as viral proteins with similar molecular weight such as NSP8 and 17 other viral proteins did not cause cell killing, nor do they induce the ROS under the same experimental conditions. It would be of interest to identify the binding partner(s) of the ORF3a in human cells.

ORF3a mutations are associated with disease progression (16) and high mortality in patients with COVID-19 (53). The Q57H mutation (16, 53) recently was found in the emerging Beta variant (54), and the Q57H variant was suspected of contributing to a surge of SARS-CoV-2 infection in Hong Kong (55). Our data show that the Q57H mutant activities are comparable to the wildtype ORF3a regarding the induction of cellular oxidative stress, innate immune responses, and apoptosis and necrosis (**Fig. 4–5**). However, our data cannot rule out the possibility that the Q57H mutant may have other effects on viral pathogenesis. For example, the Q57H mutant was predicted to bind S protein, whereas the wildtype does not (56). It would be of interest to test whether the Q57H mutant contributes to viral entry. A number of new ORF3a mutations were found to associate with the emergence of the new viral variants. It would be necessary to test the effect of those natural ORF3a variants on their interactions with host cellular stress and innate immune responses to better understand the role of ORF3a in COVID-19.

In summary, through genome-wide characterization, twelve viral proteins were identified to exert cytopathic effects. Since these viral proteins are essential for viral survival and contribute to viral pathogenesis, they could serve as therapeutic targets for future antiviral drug discovery. In accord with this, we demonstrated here that ORF3a induces apoptosis and necrosis through the activation of cellular oxidative stress and host innate immune pro-inflammatory responses. Since these ORF3a activities are linked to viral pathogenesis, disease progression and severity of COVID-19, it would be desirable to develop a fission yeast cell-based high throughput system to identify anti-ORF3a drugs to battle COVID-19.

## Materials and Methods

### Cell and Growth Media

A wild-type fission yeast SP223 strain (*h-, ade6-216, leu1-32, ura4-294)* was used to test the effect of SARS-CoV-2 viral proteins in this study (5, 13). Standard YES complete or minimal EMM or PMG selective media supplemented with adenine, uracil, leucine or thiamine (20 μM) was used to grow fission yeast cells or to select for plasmid-carrying cells. Luria-Bertani (LB) medium supplemented with ampicillin (100 μg/mL) was used for growing NEB Stable *E. coli* (NEB Cat#: C3040H) or DH5α cells and for DNA transformation.

Human pulmonary epithelial cell lines A549 (ATCC#: CCL-185), Calu-3 (ATCC# HTB-55), and a human embryonic kidney epithelial 293T cell line was used in this study. A549 and 293T cell lines were maintained in the high glucose Dulbecco’s modified Eagle’s medium (DMEM) (Corning Cat#: 10-017-CV) with 10% fetal bovine serum (FBS, Gibco Cat#: 100-438-026) and 100 U/mL Penicillin-Streptomycin (Gibco Cat#: 15140122). Calu-3 cells were maintained in Eagle’s Minimum Essential medium (EMEM) (Quality Biological Cat#: 112-018-101) with 10% FBS and 100 U/mL Penicillin-Streptomycin. All Cell lines were grown in an incubator at 37°C with 5% CO_2_.

### SARS-CoV-2 Reference Strain and Plasmids

A US SARS-CoV-2 reference strain USA-WA1/2020 (GenBank#: MN985325) was used for the genome-wide functional analysis of the SARS-CoV-2 viral proteins. A fission yeast gene cloning system, which includes a pYZ1N gene expression plasmid, has been described previously (7, 12). pYZ1N carries an inducible *nmt1* (<u>n</u>o <u>m</u>essage in the <u>t</u>hiamine) promoter. Through regulation of this promoter, the viral gene expression can be either induced or repressed in the absence (*gene*-off) or presence (*gene*-on) of 20 μM thiamine, respectively (12). The fission yeast strain that carries the pYZ1N-SARS-CoV-2 ORF was maintained in the minimal EMM medium with the selection of the *LEU2* gene carried on the plasmid. For the mammalian ORF3a study, a lentiviral constitutive expression vector pLVX-EF1alpha-IRES-Puro (Takara) that carries the ORF insert (provided by Dr. Nevan J. Krogan of UCSF) was used (3). The ORF3a mutant variants (Q57H and ΔG188) were generated by overlapping PCR with mutant-specific primers (**Table S1**) and cloned onto the same pLVX-EF1alpha-IRES-Puro plasmid *via* Gibson assembly method. All final constructs were verified by Sanger sequencing.

### Molecular Cloning of SARS-COV-2 ORFs in Fission Yeast

A total of 29 SARS-CoV-2 viral ORFs were cloned into a fission yeast pYZ1N gene expression vector system using methods as described previously (5, 7) (**Fig. S1-a**). Among them, 27 of the viral ORFs were described previously (3). NSP3 or NSP16 was obtained from Zhe Han of University of Maryland, Baltimore, or Fritz Roth of University of Toronto (Addgene plasmid # 141269), respectively. Briefly, each SARS-CoV-2 protein-encoding nucleotide was PCR amplified with a pair of primers that contained specific restriction enzymes (**Table S1**) for molecular cloning and cloned into the pYZ1N expression vector for the functional analysis. The SARS-CoV-2 viral gene inserts were verified by restriction digestions and Sanger sequencing.

### Fission Yeast Plasmid Transformation and Inducible SARS-COV-2 Gene Expression

The SARS-COV-2 gene-carrying pYZ1N plasmids were transformed into a wild type fission yeast SP223 strain by electroporation (5, 57). The plasmid transformants were selected for the presence of the *Leu*2 gene on a minimal selective PMG medium. Successful transformation of the respective SARS-COV-2-containing plasmids was verified by colony PCR with gene-specific primers (**Table S1**). Successful expression of the respective SARS-COV-2 *ORF* mRNA transcripts was measured by reverse transcription PCR using gene-specific primers (**Table S1**) and SuperScript III One-Step RT-PCR system (Invitrogen; Cat#: 12574-030). To measure SARS-COV-2 gene-specific activities, the PCR-confirmed yeast colony, which carries the desired and specific SARS-COV-2 gene-containing plasmid, was grown to log phase in liquid EMM medium supplemented with 20 μM of thiamine. Cells were then washed three times with distilled water to remove thiamine. Finally, 2 x 10^5^ cells/mL, which was quantified using a TC20 automated cell counter (Bio-Rad), were re-inoculated into fresh EMM liquid medium without thiamine to induce gene expression (*gene*-on) or with thiamine to suppress gene expression (*gene*-off). The cell cultures were grown at 30°C with constant shaking before the observation.

### Measurement of Fission Yeast Cell-specific Activities

The effects of a SARS-COV-2 protein on fission yeast cellular growth were measured by several methods, which include a yeast colony-forming assay to measure cell proliferation, a cell viability assay (5, 11), and the cellular growth kinetics to quantify cellular growth (5, 13). Briefly, the fission yeast cultures were prepared as described above. 50 μL of liquid cultures with approximately 1 × 10^3^ cells were spread onto the selective EMM agar plates with (*gene*-off) and without (*gene*-on) thiamine. The agar plates were incubated at 30°C for 4–6 days before the observation for the presence or absence of fission yeast colony formations and the sizes of the forming colonies. The absence of colonies on the agar plates indicates complete inhibition of cellular proliferation. Fewer colonies with smaller colony sizes than the normal control typically suggest reduced cellular proliferation.

To quantify the level of growth inhibition caused by the expression of a SARS-CoV-2 protein in fission yeast, 100 μL of *gene*-on or *gene*-off liquid cultures (2 × 10^5^ cells/mL) were grown in the 96-well microtiter plate in the selective EMM medium over time at 30°C in an incubator with moisture. Cellular growth kinetics was measured at OD_650_ over 48 hours by using a Synergy™ H1M monochromator-based multimode microplate reader (BioTek).

The methods used to test the effect of each viral protein production on fission yeast cell morphology have been described previously (5, 13). Briefly, fission yeast cell morphology was observed using a BZX fluorescence microscopy (Keyence) under the bright field 48 h *agi*. The overall cell morphology was evaluated by flow cytometry using forward scattered analysis (5, 13). Ten thousand cells were analyzed on a FACSCanto II flow cytometer (Becton Dickinson). The forward-scattered light (FSC) and side-scattered light (SSC) were measured for each cell population. FSC is proportional to the cell surface area and thus measures cell size. SSC determines intracellular complexity because it is proportional to cell granularity.

SARS-CoV-2 protein-induced cell death was detected by the Trypan blue staining (5). Trypan blue is a dye that specifically detects dead cells. The activation of fission yeast cellular oxidative stress by SARS-COV-2 proteins was detected by the production of ROS, which can be detected by a ROS-specific dye, DHE (Sigma). DHE generates red fluorescence in the presence of ROS (5, 58). DHE was added in a final concentration of 5 μg/mL. Cellular oxidative stress was measured 48 h *agi*.

### Measurement of Mammalian Cell-specific Activities

2 x 10^4^ 293T or 1 x 10^4^ A549 and Calu-3 cells/well were seeded into a 96-well plate and cultured at 37°C/5% CO2 overnight. Plasmid was transfected into cells using the Lipofactamine 3000 reagent (ThermoFisher) following the manufacture’s protocol. Cellular growth or cell death at the indicated time was quantified by cell number counting and trypan blue staining. Briefly, cells were trypsinized. 10 μL of cell suspension was mixed with equal volume of trypan blue. Within 5 minutes of mixing, the total cells and dead cells were counted using a TC20 automated Cell Counter. Cell viability was determined by the MTT assay as described (59). At the time of the measurement, 10 μL of 5 mg/mL MTT was added to each well of the 96-well plate and incubated at 37°C for 2–5 h. After the media was removed, 100 μL of DMSO was added to each well and mixed by pipetting. The plates were gently agitated for 15 minutes and the absorbance was measured at 570 nm using the H1M microplate reader.

Cell apoptosis and necrosis were measured by a RealTime-Glo™ Annexin V Apoptosis and Necrosis Assay (Promega) (60). At 24 h post plasmid transfection, the apoptosis and necrosis detection reagents were added into each well of 96-well plate. Following incubation at 37°C/5% CO2 at the indicated time, luminescence (RLU) and fluorescence (RFU, 485nmEx/520–30nmEm) were used to measure apoptosis and necrosis, respectively with the H1M microplate reader.

The activation of cellular oxidative stress in mammalian cells was detected using a ROS detection cell-based assay kit (Cayman Chemical; Cat#: 601290). The fluorescence of DHE staining was visualized using a BZX fluorescent microscope (Keyence). The intensity was measured using the H1M microplate reader.

Western blot analysis was carried out as described previously (59). Total proteins were extracted from transfected 293T cells using RIPA lysis buffer (Millipore Sigma) at 48 *hpt*. Equal amounts of total protein were separated on SDS-PAGE gel by electrophoresis and transferred to a polyvinylidene difluoride (PVDF) membrane (Bio-Rad). Target proteins were detected using the following antibodies: mouse anti-Strep tag II (Millipore Sigma: 71590); rabbit anti-Caspase 3 (Cell signaling: 9662); rabbit anti-Cleaved-Caspase 3 (Cell signaling: 9664) and mouse anti-GAPDH antibody (Cell signaling: 2118).

Real-time quantitative RT-qPCR (qRT-PCR) was carried out as we described previously (60). Briefly, a total of 1 μg of extracted total RNA was used for synthesis of first strand cDNA using a High-capacity RNA-to-cDNA kit (Thermo Fisher Scientific) according to the manufacturer’s instructions. Real-time PCR was performed on a QuantStudio 3 Real-time PCR system using gene-specific primers (**Table S2**) and 2x SYBR Green qPCR Master Mix (Bimake). The amplification conditions were 40 cycles of 95°C for 10 s and 60°C for 30 s, followed by melting curve analysis. Fold-change in mRNA expression was quantified by calculating the 2^-ΔΔCT^ value, with glyceraldehyde-3-phosphate dehydrogenase (GAPDH) mRNA as an endogenous control.

### Statistical Analysis

Pair-wise t-test or one-way ANOVA was calculated using software Prism 9 (GraphPad, San Diego, CA, USA). Statistical significance was accepted at the 95% confidence level (p < 0.05). * or #, p < 0.05; ** or ##, p < 0.01; *** or ###, p < 0.001.

## Acknowledgments

The authors would like to thank Dr. Nevan J. Krogan of UCSF (3), Dr. Zhe Han of UMB and Dr. Fritz Roth of University of Toronto (Addgene plasmid # 141269) for providing the SARS-CoV-2 plasmids used in this study. This study was supported in part by grants from NIH R21 AI129369, NIH R01 GM127212/AI150459, VA I01BX004652 and an intramural funding from the University of Maryland Medical Center (R.Y.Z.). J.M.S. is supported by grants from the Department of Veterans Affairs (I01RX003060; 1I01BX004652), the Department of Defense (SC170199), the National Heart, Lung and Blood Institute (R01HL082517) and the NINDS (R01NS102589; R01NS105633). V.G. is supported by NIH R01NS107262. Q.T. is supported by NIH/NIAID and G12MD007597. Author contributions: R.Y.Z. and J.T.Z. designed experiments and wrote the manuscript; J.T.Z., Q.L., R.S.C.C. and V.G. carried or assisted the experiments. J.M.S., Q.Y.T. and J.T.Z. participated in the writing and revision of the manuscript.

## Figure Legend for Supplementary Materials

**Figure S1.**
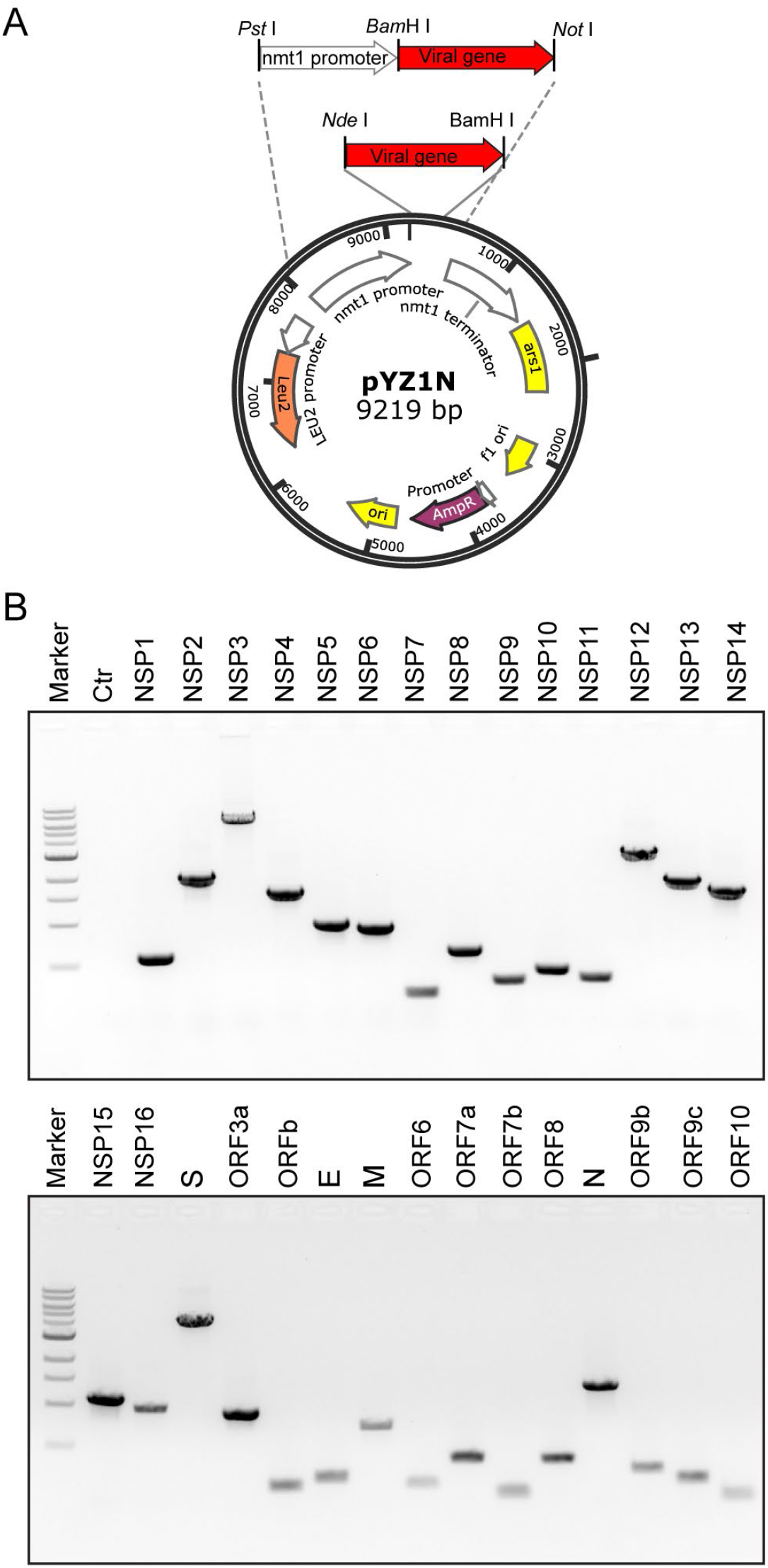
Confirmation of molecular cloning of the SARS-CoV-2 viral genome into the fission yeast gene expression pYZ1N system. (**A**) Map of the pYZ1N gene expression vector and cloning sites. (**B**) Results of RT-PCR amplified products of each SARS-CoV-2 ORF-encoding DNA sequences from SARS-CoV-2 viral cDNA.

**Figure S2.**
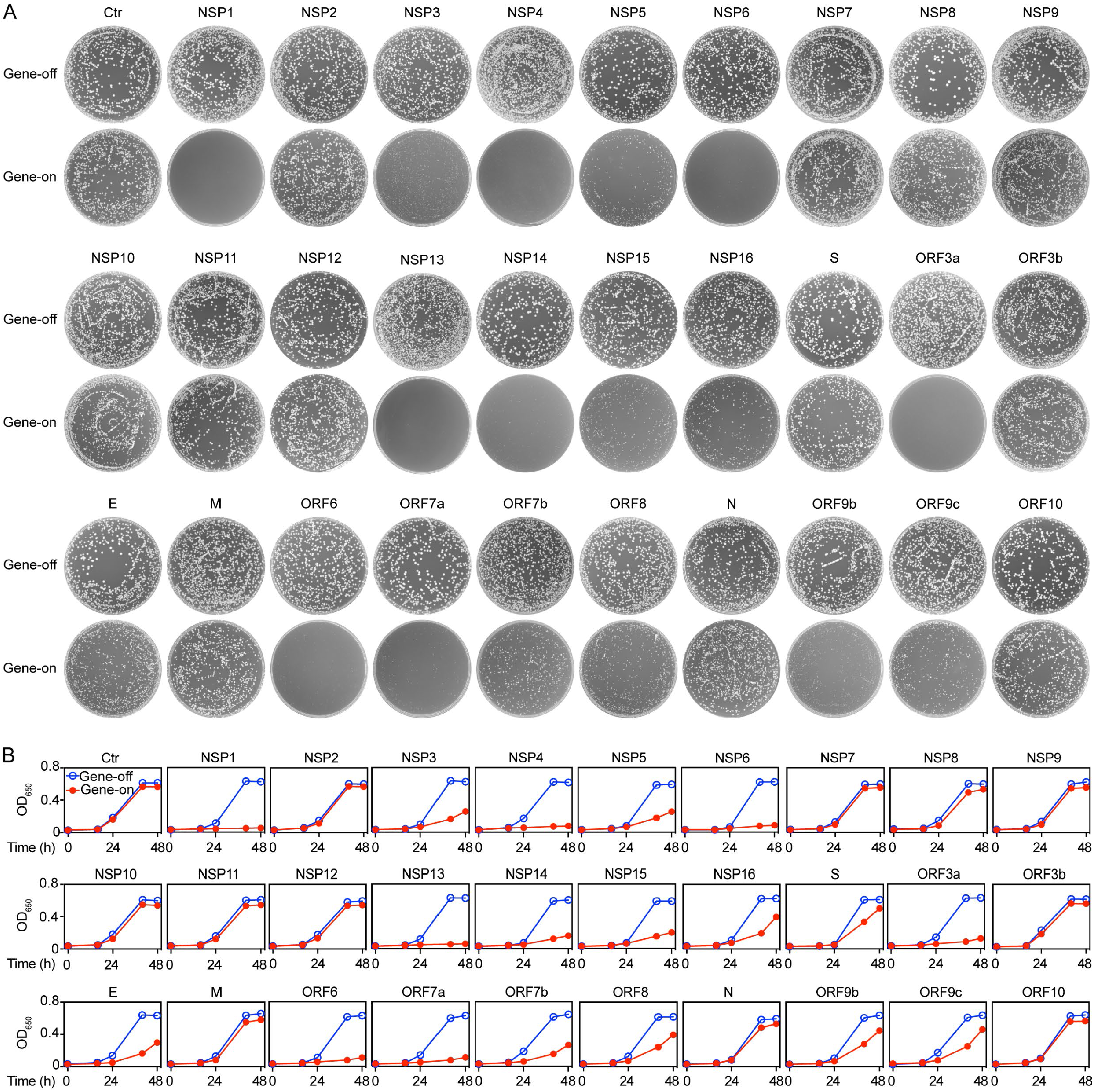
The effect of SARS-CoV-2 protein on cell proliferation. Each SARS-CoV-2-encoding ORF was cloned from the viral genome of the USA-WA1/2020 viral strain (GenBank#: MN985325) (61). Effects of SARS-CoV-2 protein production on fission yeast colony formation (**A**) and cell proliferation (**B**). Fission yeast colony formation was measured by growing SARS-CoV-2 protein-expressing fission yeast cells on the selective EMM agar plates and incubated at 30°C for 3–5 days before the pictures were taken. Cell proliferation analysis was carried out by comparing cellular growth between the SARS-CoV-2 protein-expressing cells and the SARS-CoV-2 protein-suppressing cells over time. Cell growth was measured by OD_650_ using a spectrophotometer. Ctr, empty vector control; Gene-off, no SARS-CoV-2 protein production; gene-on, SARS-CoV-2 protein produced.

**Figure S3.**
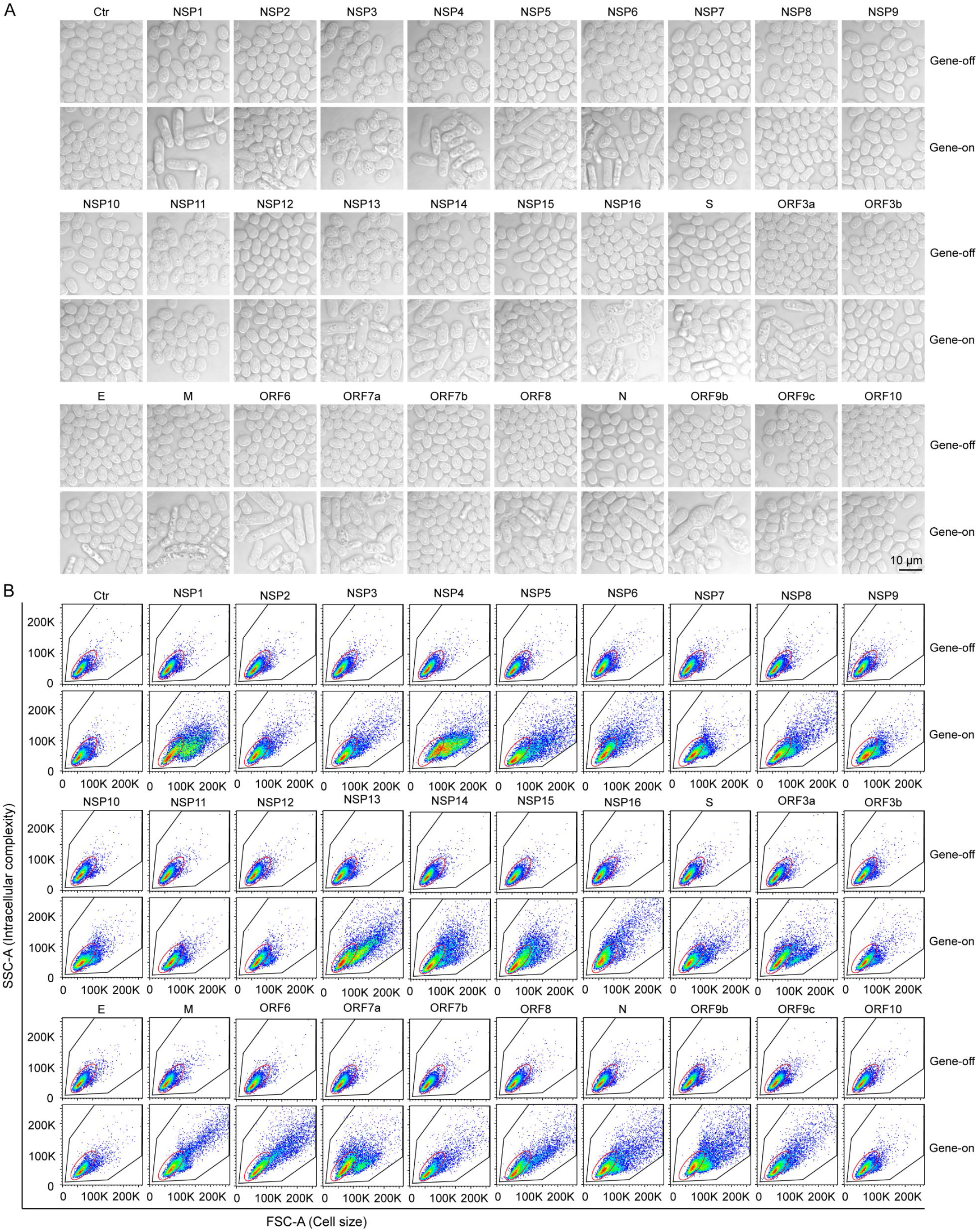
SARS-CoV-2 protein-induced cell morphologic changes in fission yeast. (**A**) Flow cytometric graph of cell morphology of 10,000 cells. Each image was obtained 48 h *agi* using bright field microscopy. Scale bar = 10 μM. (**B**) Overall cell morphology as shown by the forward scattered analysis. Ten thousand cells were measured 48 h *agi*. The forward-scatter (FSC) measures the distribution of all cell sizes. The side-scatter (SSC) determines intracellular complexity. Gene-off, no SARS-CoV-2 protein production; gene-on, SARS-CoV-2 protein produced.

**Figure S4.**
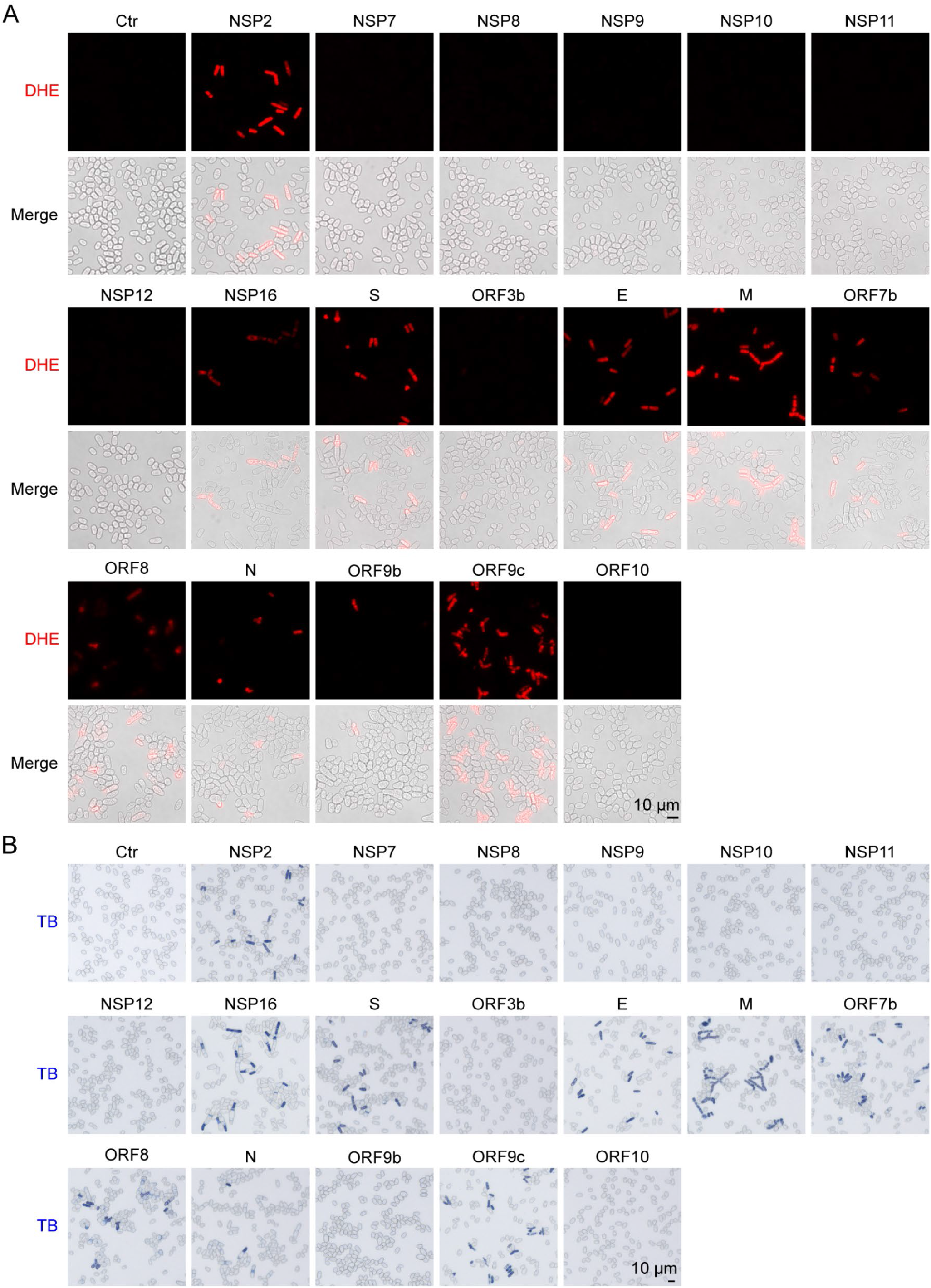
Measurement of SARS-CoV-2 protein-induced cell death and induction of cellular oxidative stress. (**A**) Measurement of SARS-CoV-2 protein-induced oxidative stress by the DHE staining showing the presence or absence of the ROS production. Images were taken 48 h *agi*. BF, bright field; (**B**) Measurement of SARS-CoV-2 protein-induced cell death 48 h *agi* by the trypan blue staining. The SARS-CoV-2 cytopathic proteins that are shown in **Figure 3** are not included here. Note that a number of other proteins also induce ROS production and cell death but at levels less than those cytopathic proteins shown in **Figure 3**. Those proteins include structural proteins M, S and E proteins, NSP2 and ORF9c that are not discussed in this manuscript. Scale bar = 10 μM.

**Table S1.**
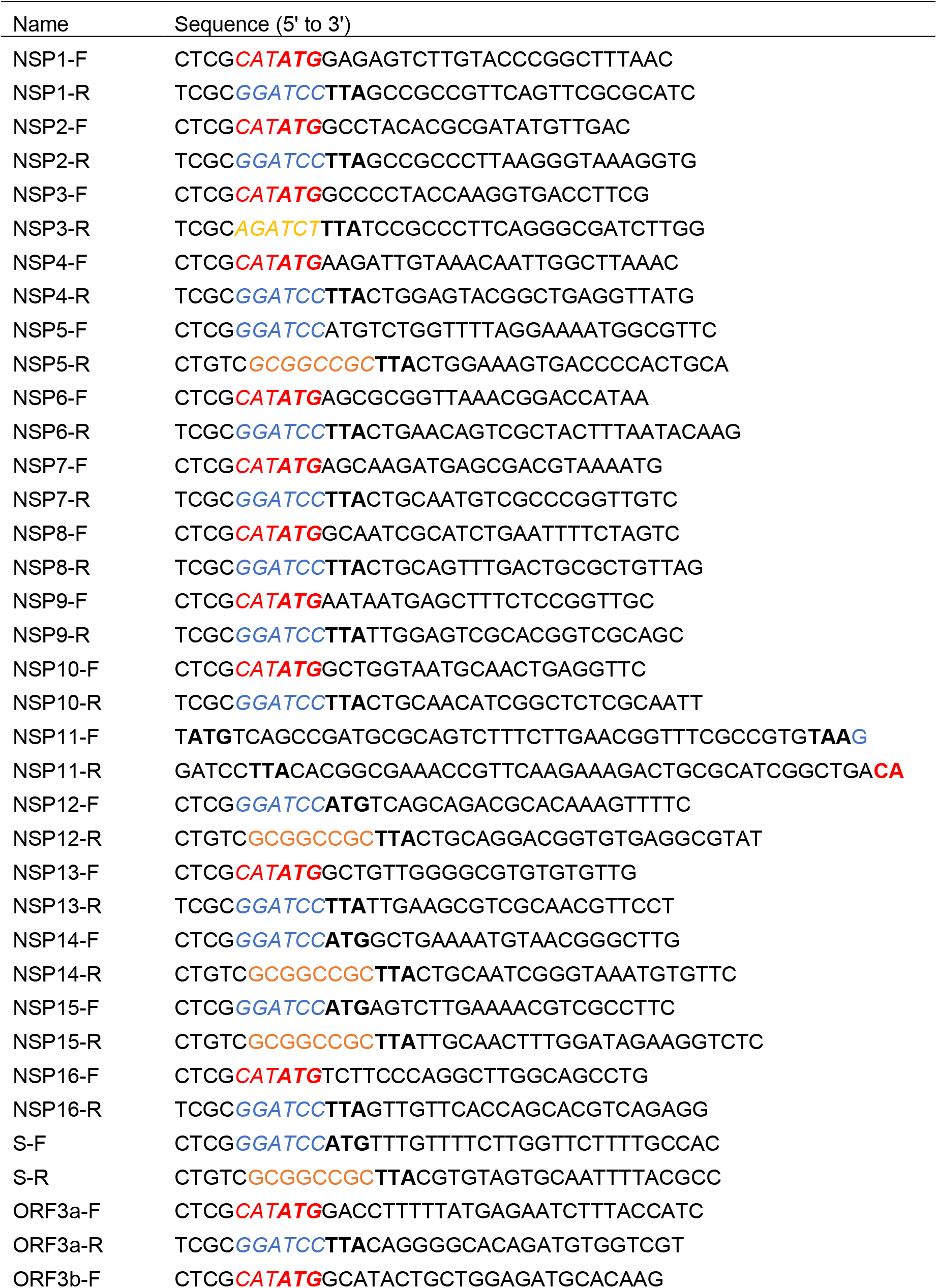
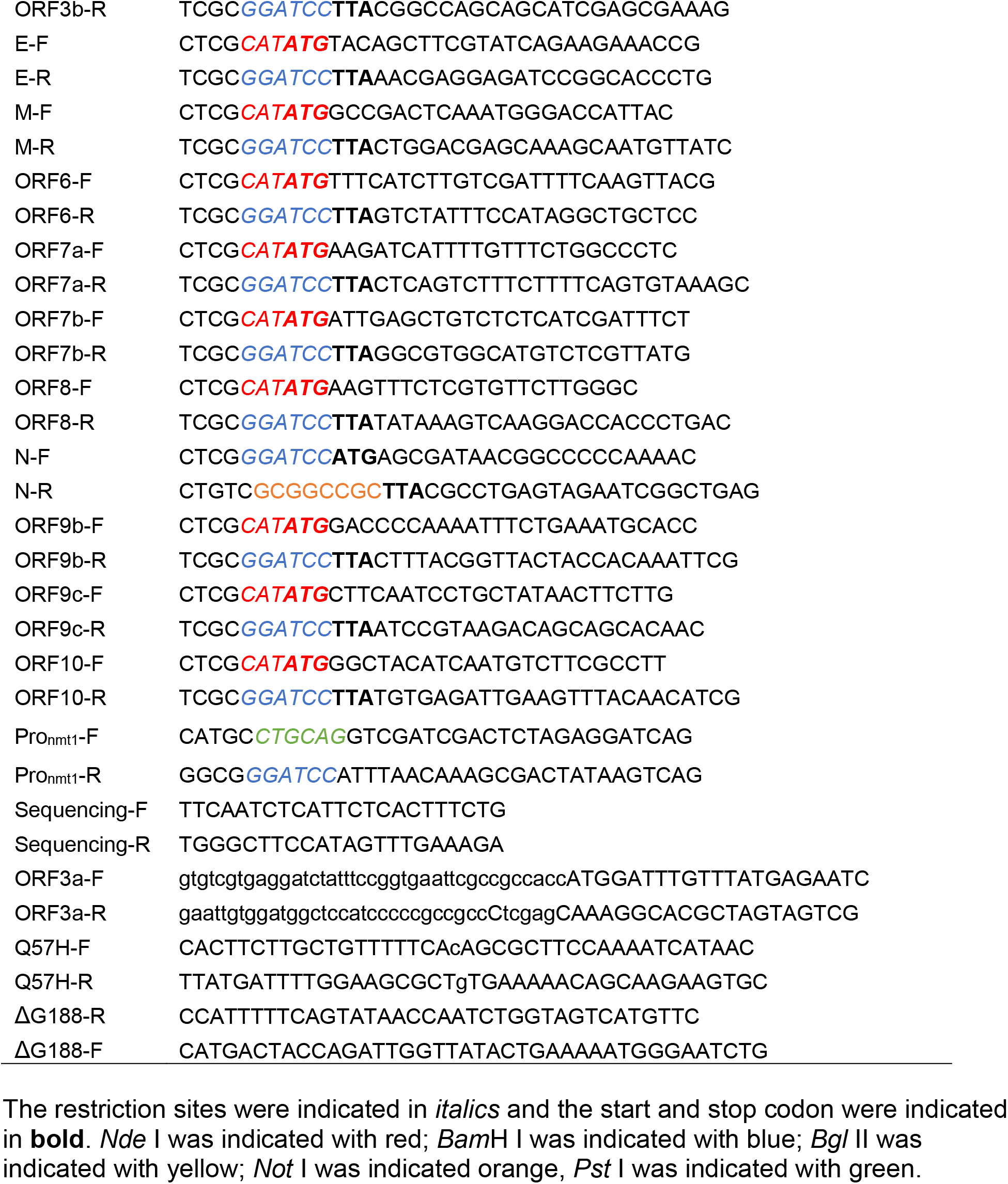
Primers used for gene cloning in this study.

**Table S2.**
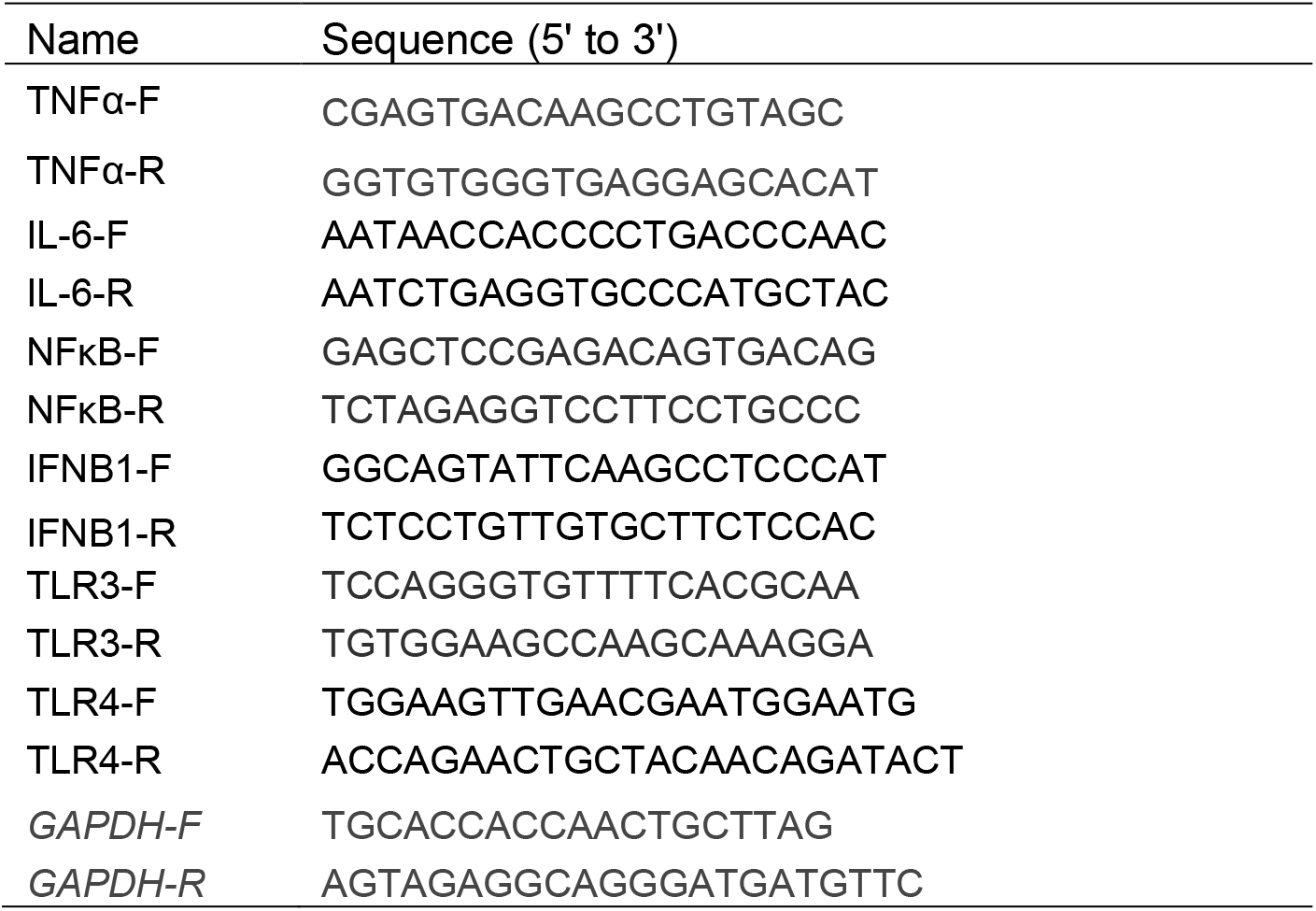
Primers used for real-time PCR in this study.

